# Thyroid hormone receptor beta (THRB) dependent regulation of diurnal hepatic lipid metabolism in adult male mice

**DOI:** 10.1101/2024.01.30.577730

**Authors:** Leonardo Vinicius Monteiro de Assis, Lisbeth Harder, Julica Inderhees, Olaf Jöhren, Jens Mittag, Henrik Oster

**Author notes:** Corresponding author: Leonardo VM de Assis, Center of Brain Behavior & Metabolism, Institute of Neurobiology, University of Lübeck, Germany, Marie Curie Street, 23562 Lübeck, Germany. or.

## Abstract

Thyroid hormones (THs) are critical regulators of systemic energy metabolism and homeostasis. In the liver, high TH action protects against steatosis by enhancing cholesterol and triglyceride turnover, with thyroid hormone receptor beta (THRB) signaling playing a pivotal role. This study probed the potential interaction between THRB action and another critical regulator of liver energy metabolism, the circadian clock. Liver transcriptome analysis of THRB deficient (THRB^KO^) mice under normal chow conditions revealed a markedly modest impact of THRB deletion. Temporal transcriptome and lipidome profiling uncovered significant alterations in diurnal metabolic rhythms attributable to THRB deficiency pointing to a pro-steatotic state with elevated levels of cholesterol, tri- and diacylglycerides, and fatty acids. These findings were confirmed by THRB agonization in hepatocytes under steatosis-promoting conditions *in vitro*. Integration of transcriptome profiles from THRB^KO^ mice and mice with induced high or low TH action identified a subset of TH responsive but THRB insensitive genes implicated in immune processes. In summary, our study reveals a complex time-of-day dependent interaction of different TH-related signals in the regulation of liver physiology indicating an opportunity for chronopharmacological approaches to TH/THR(B) manipulation in fatty liver diseases.

## INTRODUCTION

Thyroid hormones (THs) – thyroxine (T_4_) and its main biologically active derivative, triiodothyronine (T_3_) – regulate systemic energy homeostasis through transcriptional programs in metabolic tissues. TH action in target tissues mainly stems from T_3_ binding to nuclear hormone receptors alpha and beta (THRA and THRB – the latter being the predominantly expressed form in liver). Upon T_3_ binding, transcriptional co-factors are recruited and target gene transcription is initiated ^1–4^. In the liver, THs stimulate lipogenesis and triglyceride storage, lipid breakdown by boosting β-oxidation, and ATP generation by targeting β-oxidation-derived acetyl-CoA to the TCA cycle. THs further modulate cholesterol metabolism by augmenting its biosynthesis, serum uptake, and conversion into bile acids. Overall, an elevated liver TH state results in reduced fatty acid and cholesterol levels, thus making it an attractive pharmacological target for the treatment of metabolic dysfunction associated steatotic liver disease (MASLD) or non-alcoholic fatty liver disease (NAFLD) ^5,6^.

Temporal regulation of physiological functions is vital for an organism’s adaptation to cyclic environmental changes. Animals have developed a so-called circadian timing system capable of detecting and predicting external daytime and adjusting physiological processes accordingly. At the molecular level, the circadian system is based on a set of interlocked transcriptional and translational feedback loops that drive rhythmic expression of tissue-specific clock target genes ^7,8^.

Many endocrine signals such as cortisol or melatonin exhibit marked circadian secretion patterns ^9,10^. However, to which extent THs are subject to circadian control is still debated. Previous studies show that blood T_3_ and T_4_ levels show no or only weak circadian variations. At the same time, however, substantial circadian alterations in systemic energy state, and in hepatic transcriptome regulation – including TH target genes involved in fatty acid and cholesterol metabolism – have been observed. Interestingly, changes in TH state have only minor effects on the regulation of the hepatic circadian clock machinery ^11^.

In this study, we investigated the role of THRB as the primary liver TH receptor isoform in regulating diurnal transcriptome and lipidome rhythms in this organ. Our findings indicate an interplay between the circadian clock and hepatic TH action. Loss of THRB led to a combination of baseline and phasing effects in the transcriptional signature of metabolic processes indicative of a pro-steatotic state. In line with this, lack of THRB significantly impacted the diurnal lipidome of the liver with a net-increase in fatty acid and cholesterol levels. *In vitro* agonization of THRB in steatotic hepatocytes confirmed this role of THRB action as therapeutic target for MASLD.

## RESULTS

### Loss of THRB has modest effects on liver transcriptome state

To evaluate the consequence of impaired THRB signaling on liver physiology, we used a knockout model in which the DNA binding activity of THRB is impaired ^12^. Adult male wild-type (THRB^WT^) and *Thrb* knockout mice (THRB^KO^) were kept under a standard 24-hour light:dark cycle and received a regular chow diet and water *ad libitum*. Mice were culled at 4-hour intervals for 24h, and liver samples were subjected to RNA sequencing. To achieve a time-of-day independent comparison of transcriptome state between both genotypes, all sampling points were pooled (Figure 1A, upper panel). A total of 99 differently expressed genes (DEGs; up n = 50; down n = 49) were identified in THRB^KO^ livers including the TH modulator *Dio1* (down) and the lipoprotein receptor *Lrp2* (up) (Figure 1B, Table S1). Compared against DEGs from previous studies in mice with systemically low (methimazole plus potassium perchlorate treatment; MMI^13^) or high (0.5 mg/mL supplementation of T_3_ ^11^) TH activity, the extent of regulation in THRB^KO^ mice was relatively low suggesting THRB independent effects of TH action on liver transcription (Figure 1C).

**Figure 1:**
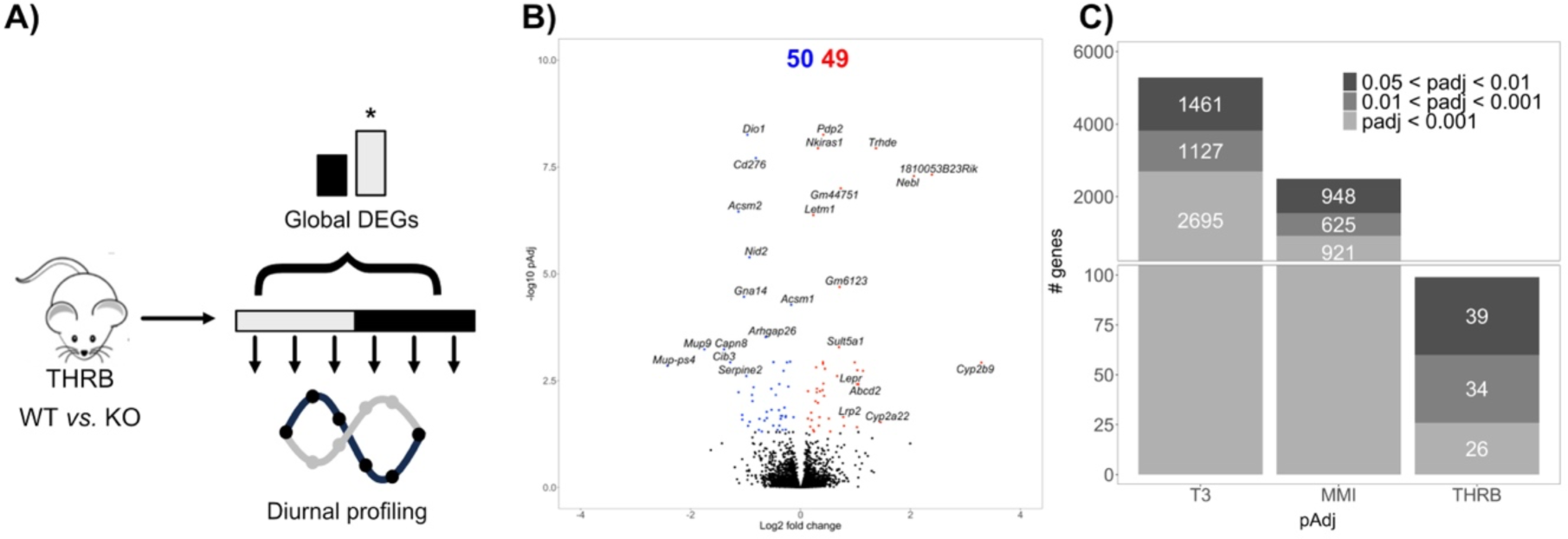
Deletion of THRB has only modest overall effects on the liver transcriptome. A) Experimental design and modes of DEG analysis. B) Volcano plot of global differentially expressed genes (i.e., DEGs irrespective of sampling time; padj < 0.05). Red and blue represent significantly UP- and DOWN-regulated genes, respectively. C) Comparison of global DEG numbers between livers from T_3_- and MMI-treated and THRB-deficient animals stratified for significance threshold. N = 3 independent samples per time point and condition for THRB^WT^ and THRB^KO^ mice.

### Loss of THRB alters diurnal phase of liver biological processes transcript rhythms

To get more insight into the effects of THRB deletion on liver transcription, we stratified our data for sampling times and analyzed changes in circadian rhythm parameters (Figure 1A, lower panel). A combination of complementary algorithms (JTK cycle, Metacycle, DryR, and CircaN) were used to assess circadian rhythms in transcript regulation, enhancing reliability and mitigating specific limitations inherent to each individual method ^14^. This approach identified 9,746 and 9,271 genes showing significant circadian regulation in THRB^WT^ and THRB^KO^ liver samples, respectively (Figure 2A; Table S2). Notably, peak phase distributions of these circadian genes were consistent across genotypes with lowest activity in the first half of the dark phase (ZT12-18; Figure 2B).

**Figure 2:**
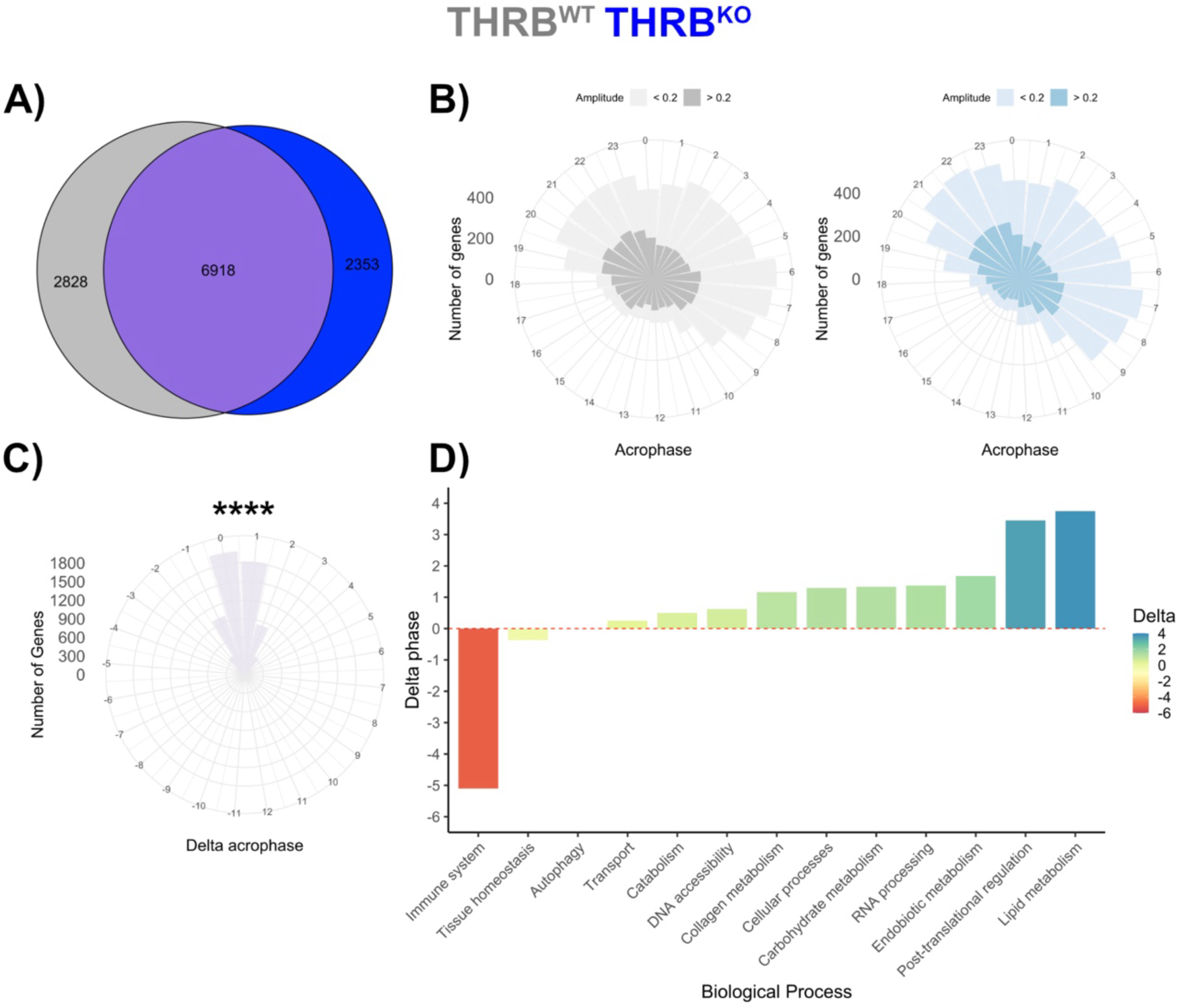
Loss of THRB temporally segregates liver immune system and metabolism-associated transcripts. A) Rhythmic genes after diurnal profiling (using JTK_cycle, Metacycle, DyrR, and CircaN) are shown in a Venn diagram for THRB^WT^ (grey) and THRB^KO^ livers (blue). B – C) Acrophase distribution of rhythmic transcripts in livers of THRB^WT^ and THRB^KO^ livers (B) and phase difference for robustly rhythmic genes between both genotypes (C). D) THRB-dependent acrophase shifts of rhythmic transcripts of different physiological processes are depicted. ****: p < 0.0001. N = 3 independent samples per time point and condition.

For those genes with significant rhythmicity in both genotypes, we observed a slight but significant phase delay of 0.4 hours in THRB^KO^ compared to THRB^WT^ livers (Figure 2 B– C). Using the phase set enrichment analysis (PSEA) algorithm ^15^, we further assessed the impact of THRB on the timing of specific biological processes. Similar processes were manually combined into broader biological categories to streamline the data. This approach yielded a general delay of up to 3.5 h in expression rhythms associated with metabolism while immune system-related processes showed a marked delay of around 5 h in THRB^KO^ mice (Figure 2D; Table S2).

### Loss of THRB alters diurnal rhythms of fatty acid and cholesterol metabolism

For a more detailed analysis of circadian deregulation in THRB-deficient livers, we directly compared transcript rhythm parameters between both genotypes. CircaCompare ^16^ analysis identified 1,446, 314, and 369 genes with changes in mesor (baseline expression), amplitude (time-of-day dependent variation), and acrophase (time of peak expression), respectively, termed as *differentially rhythmic genes* (DRGs, Figure 3A; Table S3). Loss of THRB affected liver transcriptome rhythms predominantly at the mesor level (764 mesor up and 682 mesor down) with slightly lower numbers for phase (98 advanced and 271 delayed genes) and amplitude (207 and 107 genes with lower and higher amplitude, respectively) (Figure 3A). Gene set enrichment analysis (GSEA) of DRG classes for biological processes revealed a complex regulation at the transcriptome level in the absence of THRB. For instance, DRGs with reduced and increased mesor enriched for mRNA catabolic processes and phosphorylation, respectively. Conversely, DRGs with reduced amplitudes enriched for AKT pathways (PI3K pathway and AKT signaling), protein ubiquitination, and response to glucose. DRGs with increased amplitude, on the other hand, were enriched for transcriptional regulation, proteolysis, and protein kinase activity (Figure 3 B; Table S3). Most strikingly, circadian profiling allowed us to identify a dual regulation at the mesor level for transcripts associated with fatty acid and cholesterol regulation. GSEA from DRGs with increased mesor effect enriched for cholesterol and bile acid metabolism (*Acnat2, Abcb11, Akr1d1, Baat, Cyp7a1*, *Npc1*, *Slc27a2, Hmgcs2, Vldlr*) as well as fatty acid metabolism and beta-oxidation (*Abcd2*, *Acox2, Acot3, Acot12, Cpt1a, Ehhadh, Slc27a2*). Conversely, mesor down DRGs enriched for acyl-coA biosynthesis, hydrolysis, and fatty acid transport (*Acsm1*, *Acsm2*, *Acaa1a*, *Abhd6, Acox3, Fabp1*, *Fabp2*, *Slc27a3*, *Slc27a4*) (Figure 3B). These analyses predicted that DRGs with increased mesor favor, both, higher fatty acid synthesis and oxidation (*Acox2*, *Acnat2, Acad11*, *Acot3, Acot12*, *Cpt1a, Ehhadh*, *Scd2*, *Slc27a2*, *Hsd17b4, Hpgd*) whereas DRGs with reduced mesor point to a decreased fatty acid breakdown and transport (*Acsm1*, *Acsm2*, *Acot6*, *Acox3*, *Abhd6*, *Fabp1*, *Fabp2*, *Mboat1, Slc27a3*, *Slc27a4*). Genes pertaining to cholesterol synthesis and bile acid secretion (*Baat*, *Akr1d1*, *Cyp7a1*, *Hmgcs2*, *Npc1*, *Vldlr*) were identified only in DRGs with increased mesor, which suggests increased cholesterol metabolization and secretion as bile acids (Figure 3 C,D). Circadian profiling of pathway associated genes confirmed the mesor effects with higher effects towards the end of the dark phase (ZT18-24) (Figure 3 E; Table S3). In summary, detailed circadian profiling predicted that THRB^KO^ livers would tend towards increased fatty acid and cholesterol levels.

**Figure 3:**
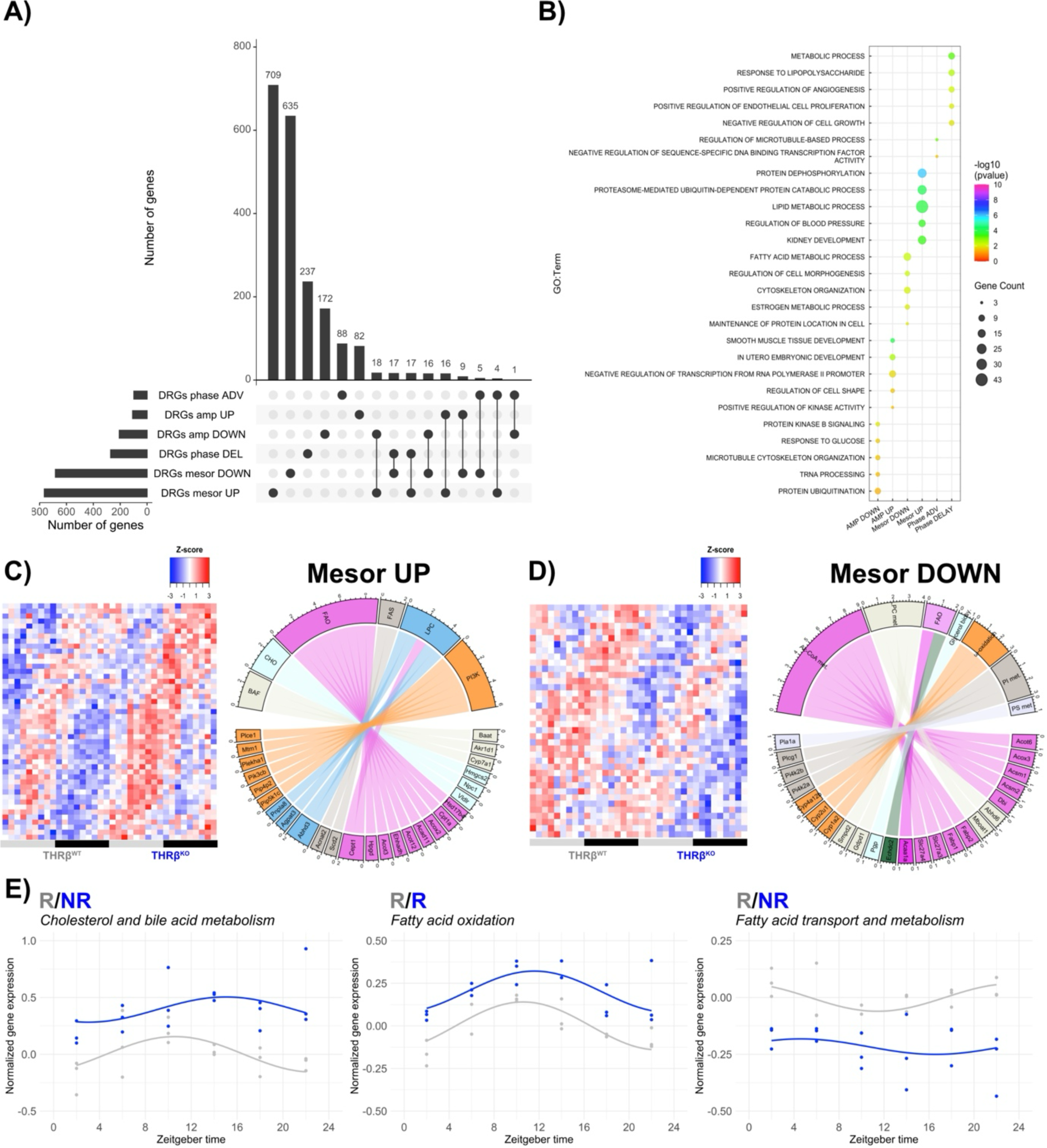
Altered diurnal rhythms in fatty acid and cholesterol metabolism in the liver of THRB^KO^ mice. A) Circadian profiling of liver transcripts (using CircaCompare) was performed, and numbers of differentially rhythmic genes (DRGs) are represented as UpSet plot. B) Gene set enrichment (GSEA) top-5 processes for each DRG class (amplitude, phase, mesor) are shown. C – D) Manual curation of lipid metabolism-associated DRGs for mesor was performed and shown as heat maps. Chord plots show gene names and associated biological processes. E) Genes associated with the biological processes were averaged and plotted across 24h and circadian analysis was performed using CircaCompare. BAF = bile acid formation. CHO = cholesterol metabolism. FAO = fatty acid oxidation. FAS = fatty acid synthesis. LPC = lysophosphatidylcholine. PI3K = phosphoinositide 3-kinase. PI and PS = phosphatidyl and phosphatidylserine metabolism. N = 3 independent samples per time point and condition. R = presence of rhythmicity. NR = absence of rhythmicity.

### Loss of THRB alters liver lipidome rhythms

To test this prediction, we performed diurnal untargeted lipidomic profiling on liver samples from both genotypes. In total, 285 distinct lipid species were detected of which 101 and 107 were significantly rhythmic in THRB^WT^ and THRB^KO^, respectively (Figure 4 A). Assessment of lipid classes showed main genotype differences for ceramides (more rhythmic species in THRB^KO^), phosphatidylethanolamines (more), and diacyl- and triacylglycerides (less rhythmic species in THRB^KO^) (Figure 4 B). No overall phase difference between robustly rhythmic lipids was identified between the genotypes (mean = 0.15h, p = 0.60, Figure 4 C; Table S4).

**Figure 4:**
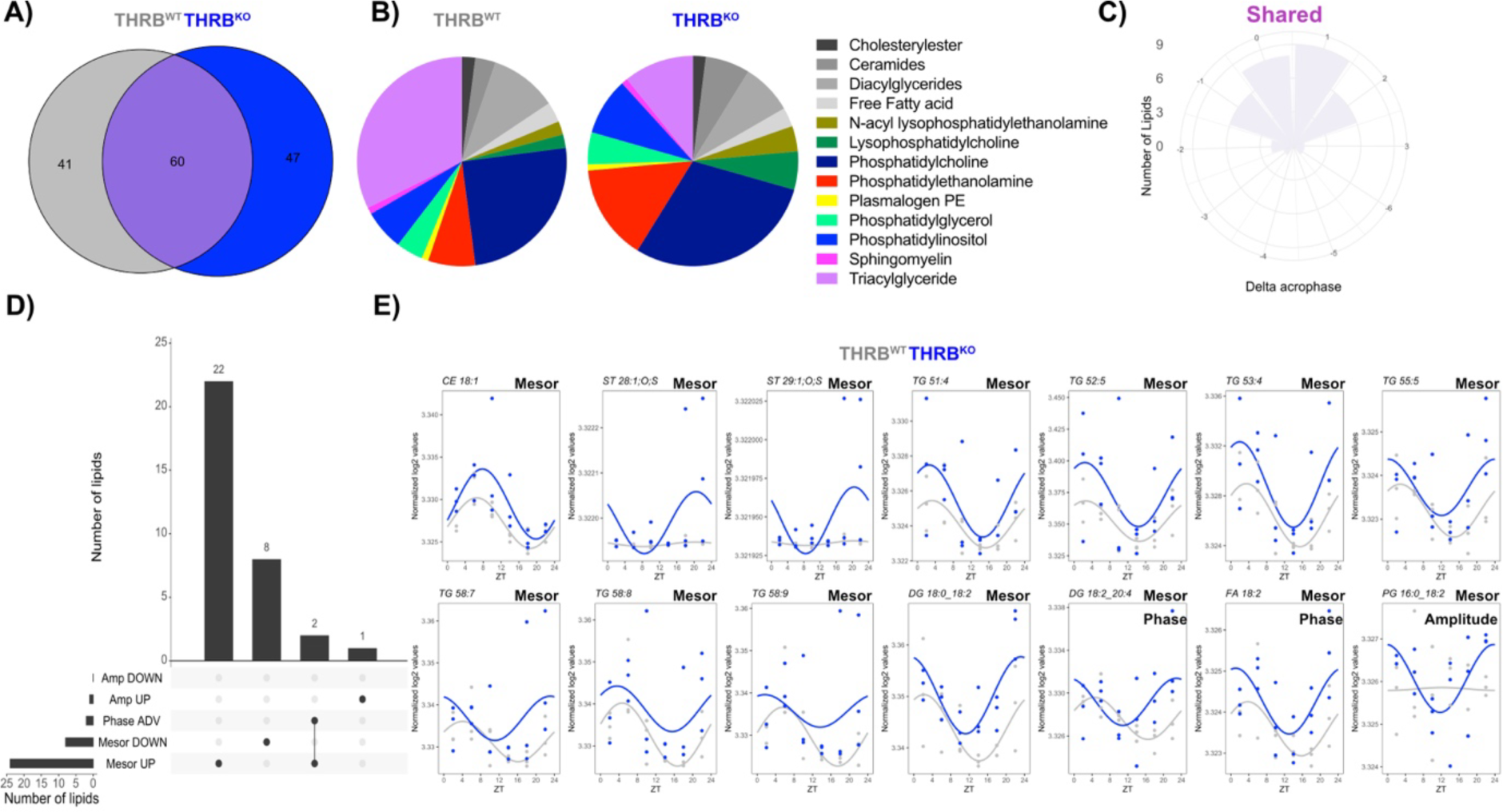
Loss of THRB alters liver lipidome rhythms with increased cholesterol, fatty acids, di- and triglycerides. A) Rhythmic lipids after diurnal profiling (using JTK_cycle, Metacycle, DyrR, and CircaN) are shown in a Venn diagram for THRB^WT^ (grey) and THRB^KO^ livers (blue). B) Distribution of lipid classes identified as rhythmic in THRB^WT^ (left) or THRB^KO^ livers. C) Rose plot depicting acrophase differences of robustly rhythmic lipids between genotypes. D) UpSet plot representing numbers of lipids with specific rhythm parameter changes (mesor, amplitude, phase) identified by CircaCompare. E) Examples of lipids with alterations in specific rhythm parameters. N = 3 independent samples per time point and condition.

Detailed comparison of lipid rhythms revealed marked changes mostly at the mesor level (Figure 4 D; Table S5). Lipid classes such as phosphatidylcholine (PC 42:2, 31:1, 18:0_22:5, 20:0_20:4, 22:0_20:4), plasmalogen PC (38:2), phosphatidylinositol (PI, 18:0_22:6, 20:0_20:4) showed mesor down effects in THRBKO. Increased mesor, in turn, was observed for cholesterol esters and sulfates (CE 18:1, ST 28:1;O;S, ST 29:1;O;S), triacylglycerols (TG 58:9, TG 58:8, TG 53:4, TG 58:7, TG 52:5, TG 55:5, TG 51:4), diacylglycerols (DG 18:1_22:6, DG 18.2_20.2, DG 18.2_20.4, DG 18.1_18.2, DG 18.0_18.2, DG 16.0_18.2), phosphatidylglycerols (PG 18:2_22:6, PG 18:1_22:6, PG 18:1_18:1), free fatty acids (18:2), ceramides (Cer 42:3;2O; Cer 18:1;2O/23:0), and phosphatidylcholines (PC 18:1_20:3, PC 18:1_18:2). Increased rhythm amplitudes were identified for phosphatidylglycerol (PG 16:0_18:2) while a phase advance of ca. 4 h (associated with a mesor increase) was found for two diacylglycerol entities, DG 18:0_18:2 and DG 18:2_20:4, with similar tendencies (p-value < 0.1) for TG 60.4 and DG 36.4 (Figure 4 E; Figure S2 – 3; Table S4).

In line with transcriptome findings, diurnal lipidomic profiling suggested slightly increased lipid contents (TAG, DAG, free fatty acids, cholesterol) in THRB^KO^ mice. It further identified additional lipid classes such as PCs, PIs, PGs, and ceramides whose mesor levels were also affected by the absence of THRB. Our findings suggest that loss of THRB results in changes in metabolic rhythms yielding a higher susceptibility to steatogenic stimuli.

### Activation of THRB reduces steatosis in hepatocytes *in vitro*

To directly assess the role of hepatic THRB action in the regulation of steatosis, we treated AML12 hepatocytes with palmitate with or without pharmacological activation of THRB. Palmitate treatment for 48 hours resulted in consistent levels of steatosis (Figure 5 A, B). Using a highly sensitive flow cytometry approach, we concurrently measured cell death, triglyceride, and cholesterol content at the individual cell level. Palmitate treatment increased cell mortality, which was significantly reversed by co-stimulation with T_3_ or the THRB agonist resmetirom. Remarkably, resmetirom reduced cell death even below baseline levels of the BSA-treated group, underscoring a potent hepatoprotective effect mediated by THRB activation (Fig. 5 B, C, left panels). As anticipated, palmitate increased, whereas T_3_ and resmetirom rescued triglyceride and cholesterol levels when compared to BSA controls (Figure 5 B, C, center and right panels). The reduction by T_3_ and resmetirom fully reverted lipid content whereas cholesterol decrease was only partial (Figure 5 C). Collectively, our findings suggest a protective role of THRB in lipid metabolism under steatotic conditions.

**Figure 5:**
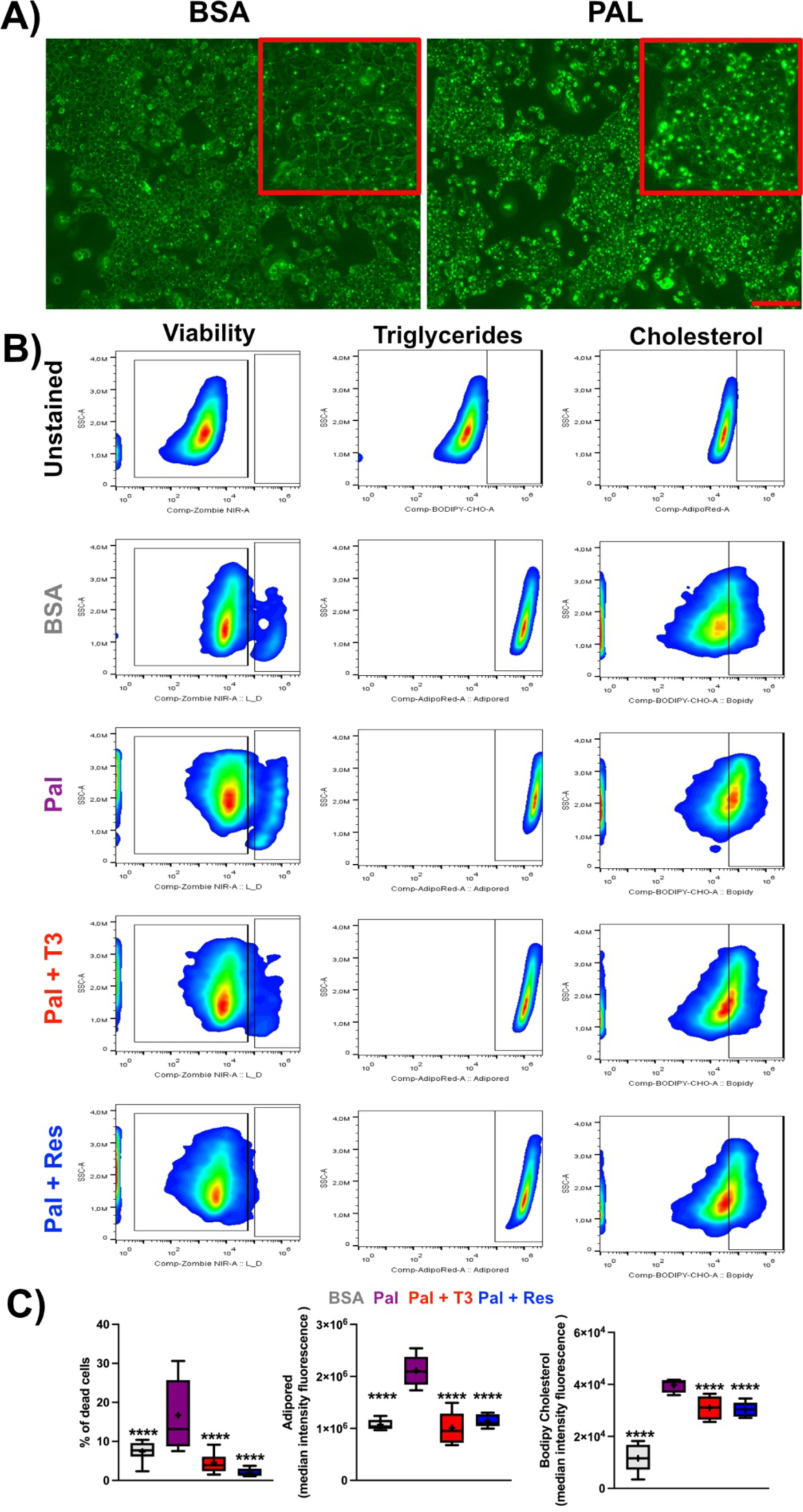
Activation of THRB reduces hepatocyte steatosis *in vitro.* A) Representative micrographs of AML-12 hepatocyte triglyceride staining (AdipoRed) under control (left) and with palmitate (right). Scale bars depict 100 µm. B) Gating strategy representing viability, triglyceride and cholesterol signal. C) Percentage of dead cells and mediant intensity fluorescence (MIF) of lipid - and cholesterol-positive cells. N = 5 – 12 independent samples per condition.

### THRB-independent TH-regulated genes are associated with immune function

Our initial findings suggested that loss of THRB has only moderate effects on overall transcriptome regulation compared to conditions with altered systemic TH state (Figure 1). THRB is recognized as the primary thyroid hormone receptor in the liver, which is largely attributed to its high prevalence in hepatocytes of the liver parenchyma ^17^. While THRA is also expressed in the liver, it has primarily been associated with other cell types ^18^. Single-cell RNA sequencing data from the Liver Cell Atlas ^19^ confirmed these findings, indicating that *Thrb* expression is mainly found in hepatocytes and Kupffer cells, while *Thra* is primarily expressed in Kupffer, T, and endothelial cells (Figure S4).

Our data suggested that many genes exhibit increased expression under high-TH conditions while being downregulated under low-TH conditions ^11,20^. Interestingly, 277 of these putative TH target genes did not show significant alterations in the absence of THRB (Figure 6A). Gene set enrichment analysis (GSEA) for these TH-regulated but THRB-insensitive genes highlighted several biological processes such as immune system, inflammation, and lipid and cholesterol metabolism (Figure 6B, Table S4). Targeted pathway analysis confirmed this notion. Interestingly, at the pathway level immune system-associated transcripts showed phase changes in line with previous PSEA data from THRB^KO^ mice (Figure 6C, 2D). These changes suggest an overall THRB-independent regulation of associated pathways by TH – possibly via THRA – which would be consistent with the predominant expression of THRA in non-hepatocyte cells.

**Figure 6:**
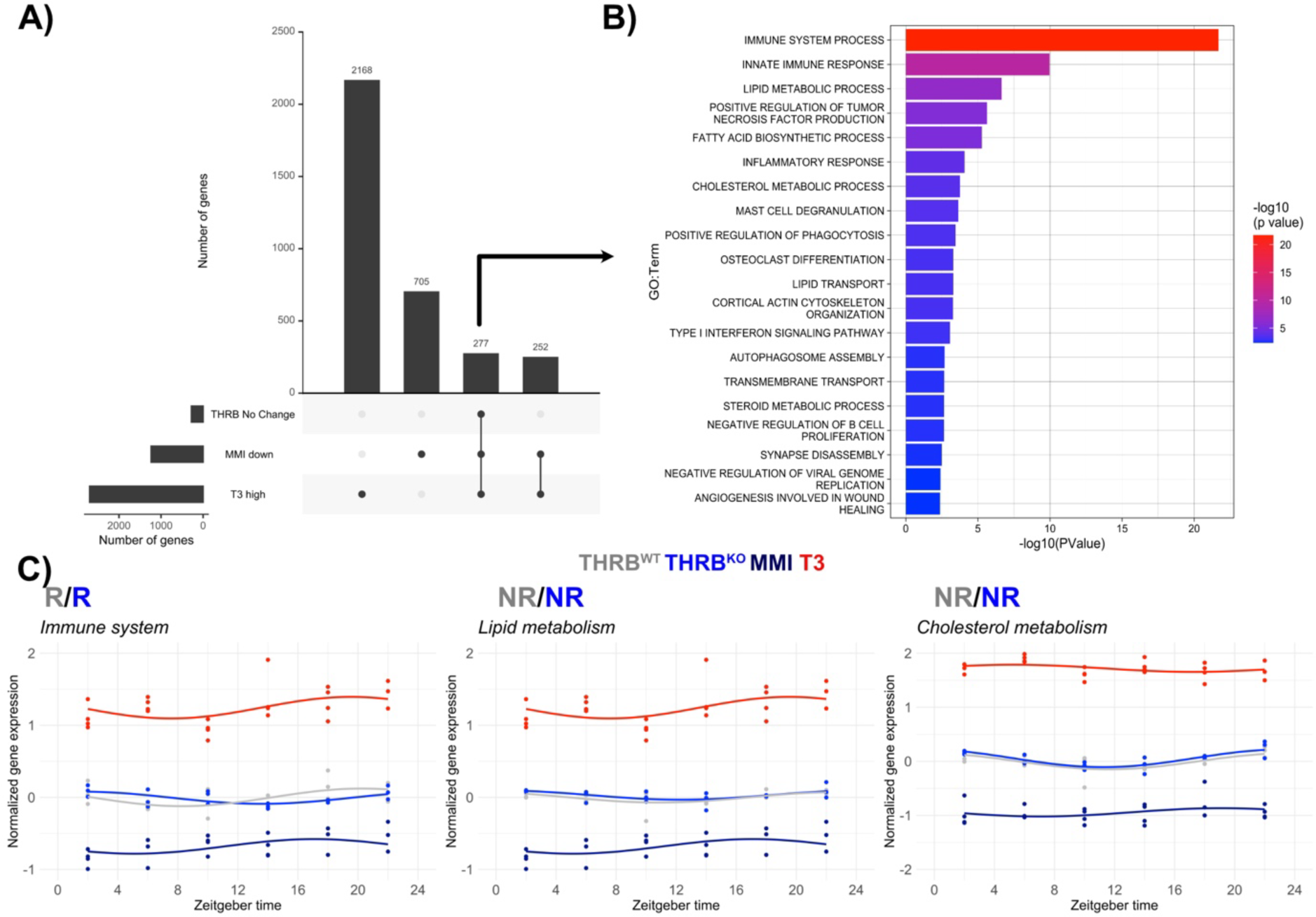
Identification and functional evaluation of THRB-independent thyroid hormone-responsive genes. A) UpSet plot comparing liver global DEGs down-regulated under low-TH (MMI) and up-regulated under high-TH (T3) conditions THRB-independent of THRB (THRB^KO^). B). Gene set enrichment analysis (GSEA) of putative THRB-independent TH targets (n = 277). C) Overall pathway expression profiles of the THRB-independent TH-responsive genes in the immune system, lipid and cholesterol metabolism in different groups. Data from low (MMI) and high (T3) levels were obtained from GSE199998. N = 3 – 4 independent samples per time point and condition.

## DISCUSSION

We show that loss of THRB results in subtle alterations in liver transcriptome regulation. Detailed circadian profile analysis unveiled alterations in the expression profiles of metabolism-associated and other genes. Particularly, in the absence of THRB, immune-related genes were predominantly phase-delayed. In contrast, genes involved in metabolic functions, such as lipid metabolism, were phase-advanced. Numerous genes associated with lipid metabolism pathways were significantly altered with regard to rhythm parameters, especially mesor. Diurnal lipidomic profiling supported these transcriptomic insights and underscored an elevation in TAGs, DAGs, and cholesterol. Cell-based assays confirm that THRB activation mitigates lipid and cholesterol build-up, underscoring its protective role against steatosis. Leveraging existing datasets, we discerned a subset of genes regulated by TH independent of THRB that are predominantly implicated in immunological and metabolic processes.

The hepatic transcriptomic response to low TH levels is much smaller compared to high-TH conditions ^13^, and this reduction is even more pronounced in THRB KO mice. This pronounced hepatic response to high TH versus the diminished response to low TH state – whether induced pharmacologically or genetically (THRB knockout) – further substantiates the notion of low hepatic TH action under physiological conditions.

We uncovered more fine-grained THRB deletion effects when considering sampling time in our analysis. Circadian profiling yielded distinct changes in transcript regulation under THRB-deficient conditions. Such an approach has previously been shown to enhance the detection of transcriptomic alterations in response to a high-fat diet ^14^. Nevertheless, the effects of THRB deletion were overall less pronounced compared to those of elevated TH conditions. Focusing on lipid metabolism, we observed bidirectional mesor level alterations: genes driving fatty acid oxidation and cholesterol metabolism were upregulated, while transcripts involved in fatty acid transport/metabolism were downregulated.

Diurnal lipidomics validated these findings, revealing accompanying changes following THRB deletion at the metabolite level. Overall, we identified 285 lipid species, with 35% exhibiting significant rhythmicity across both genotypes. This exceeds previous findings by Adamovich et al. (2014) ^21^ which reported 159 lipid entities with 17% rhythmicity. Variations in experimental protocols, such as sample processing and lipid identification, along with the methodologies for assessing rhythmicity, likely account for these differences. In a recent study, Sprenger et al. (2021) ^22^ showed that half of the liver lipidome is rhythmic, employing ANOVA for temporal analysis due to their high-resolution data and irregular sampling intervals. Direct comparisons with our results are challenging because of methodological differences. However, a consistent rhythmicity observed in TAGs across studies is notable.

33 lipids (11%) showed significant mesor variations between genotypes, and a considerable increase in cholesterol, TAGs, DAGs, and FAs in THRBKO livers was identified. We further observed significant alterations in PCs, PGs, and PIs, with most changes presenting as increased mesor levels. Notably, a specific PG species (PG 16:0_18:2) exhibited a gain of rhythmicity in THRB^KO^ mice, while some PC species showed mesor alterations (i.e., increase or decrease), thus representing a mixed effect. Similarly, specific PG and PI species were distinctly impacted in THRB^KO^ livers, displaying elevated or diminished mesor levels, respectively. Together, our findings suggest that the absence of THRB has a more pronounced impact on the liver lipidome than the transcriptome. This metabolic reprogramming lends further support to THRB’s protective role against steatosis. Lower PC/PE ratios and PC levels have been associated with metabolic syndrome-associated steatohepatitis development ^23–26^, and insulin resistance has been implicated in MASLD development ^27^ through changes in DAGs and ceramides ^28–30^. THRB KO livers show increased ceramide and TAG levels, which could predispose to MASLD development.

The ablation of THRB resulted in elevated hepatic lipid and cholesterol levels, underscoring its significant role in lipid regulation ^5,31^. Pharmacologically activating THRB has consistently been shown to decrease lipid and cholesterol levels ^17,32–34^. Contrastingly, receptor-level manipulations yielded variable results. For example, mice with a dominant-negative THRB mutation (TRBPV) display a THRB protective role, while a recent THRB knockout study did not show a preventive effect on metabolic dysfunction-associated steatohepatitis (MASH) development. In contrast, THRBKO mice were further protected against fibrosis compared to wild-type controls ^35^. These discrepancies could be due to differences in experimental approaches, such as the type of knockout and temperature conditions during experiments. Importantly, thermoneutrality-kept THRBKO mice show normalized T_3_ and T_4_ levels, whereas mice kept under non-thermoneutral conditions are hyperthyroid. It has been suggested that the availability of the thyroid hormone ligand is more critical than receptor availability in MASH. This hypothesis aligns with findings that increased hepatic TH production via upregulated DIO1 has a protective effect against MASH ^36^. Our *in vitro* experiments utilizing remestirom to assess THRB action revealed that THRB agonization reduces lipid and cholesterol levels, mimicking the hepatic action of T_3_.

THRB deletion leads to significant disruptions in lipid metabolism circadian rhythms, without concurrent changes in the expression of core circadian clock genes. This uncoupling of clock and clock outputs is reiterated in the liver under varied TH conditions, as previously shown ^11,13^, suggesting that TH may influence circadian physiology downstream of or independently from the circadian machinery. This concept remains to be further elucidated, e.g., in clock mutant models.

Our research is not without its limitations. The use of systemic THRB knockout mice introduces known non-hepatic variables, including increased serum T_3_ levels ^12^. Increased serum T_3_ could activate THRA in the liver (and other organs) and contribute to the observed effects in THRB KO mice. Additional studies with hepatocyte-specific THRB^KO^ mice and mice with deletions in the clock gene machinery are required to clarify how hepatocyte THRB regulation interacts with local clocks to shape diurnal transcriptome and lipidome regulation. Importantly, our classification of liver as a low-TH action organ is mainly based on gene expression signatures. Direct quantification of intra-hepatic TH levels is required to substantiate this concept and enzymatic profiling under different steatosis conditions to better assess the physiological consequences.

## MATERIAL AND METHODS

### Mouse model and experimental conditions

Two to three-months-old male wild type (THRB^+/−^) and knockout (THRB^−/−^) were group-housed under 12h/12h light/dark conditions (LD, 200 – 400 lux) at 22 ± 2°C with food and water provided ad libitum (normal chow, 5 % fat, 1314, Altromin, Germany). Mice were culled by cervical dislocation at 4-hour intervals, tissues were kept in dry ice, and stored at - 80°C. Animal experiments were ethically approved by the Animal Health and Care Committee of the Government of Schleswig-Holstein and in line with international guidelines for the ethical use of animals. For each timepoint, a total of 3 mice were used.

### Transcriptome analyses

Total RNA was extracted from livers using TRIzol reagent (ThermoFisher, Germany) and columns (Zymo Research, USA) according to the manufactory’s instructions. Truseq RNAseq comprising 40 million reads per sample (12GB) was performed at Novogene (Germany). Samples underwent quality control and only those with RNA integrity number (RIN) higher than 6.5 were used. Messenger RNA was purified from total RNA using poly-T oligo-attached magnetic beads. After fragmentation, the first strand cDNA was synthesized using random hexamer primers, followed by the second strand cDNA synthesis using dTTP for non-directional library. The library was checked with Qubit and real-time PCR for quantification and bioanalyzer for size distribution detection. The clustering of the index-coded samples was performed according to the manufacturer’s instructions. After cluster generation, the library preparations were sequenced on an Illumina platform and paired-end reads were generated. Raw data (raw reads) of fastq format were firstly processed through in-house perl scripts. Clean data (clean reads) were obtained by removing reads containing adapter, reads containing poly-N and low-quality reads from raw data. Q20, Q30 and GC content the clean data were calculated.

Reference genome (GRCm38/mm10) and gene model annotation files were downloaded from genome website browser (NCBI/UCSC/ENSEMBL) directly. Paired-end clean reads were mapped to the reference genome using HISAT2 software. All the downstream analyses were based on clean data with high quality. FeatureCounts was used to count the reads numbers mapped to each gene. Genes containing a sum of reads among two groups lower than 100 were excluded from the analysis. The remaining genes were log2 transformed in DESeq2 using vst function. A total of 17,994 transcripts were considered for subsequent analyses.

### Lipidomics analysis of the liver via liquid chromatography coupled to tandem mass spectrometry (LC-MS/MS)

All measurements were performed using a Dionex Ultimate 3000 RS LC-system coupled to an Orbitrap mass spectrometer (QExactive, ThermoFisher Scientific, Bremen, Germany) equipped with a heated-electrospray ionization (HESI-II) probe. All solvents were of LC-MS grade quality and were purchased from Merck (Darmstadt, Germany). Zirconium oxide beats and 500 µl of water were added to the frozen tissue (70 – 150 mg). Liver samples were homogenized for 3 min at speed level 8 (Bullet Blender, Next Advance, New York, USA) and centrifuged (2,000 g/10 min/4 °C). The supernatant (400 µl) was transferred into a new reaction tube and diluted accordingly to reach a protein concentration between 0.6 mg/ml and 1.2 mg/ml. Protein content was determined using a Bradford assay (Sigma-Aldrich, USA).

Lipid profiling was performed as described previously ^37^. Briefly, lipids were extracted by adding 1 ml of methanol/methyl tertiary-butyl ether/chloroform (1.33:1:1, v/v), containing butylated hydroxytoluene (100 mg/L, Sigma-Aldrich, Germany) and 2.5 ml/L of SPLASH internal standard mix (Avanti Polar Lipids, Alabaster, USA), to 50 µl of tissue homogenate. After incubation and centrifugation, supernatants were dried under vacuum and resuspended in 50 µl of methanol/isopropyl alcohol (1:1, v/v) for LC-MS/MS analysis. Extracted lipids were separated on an Accucore C30 RP column (150 × 2.1 mm, 2.6 µm) using acetonitrile/water 6:4 (v/v) as eluent A and isopropyl alcohol/acetonitrile (9:1, v/v) as eluent B; 10 mM ammonium acetate (Sigma-Aldrich, Germany) and 0.1 % formic acid (Biosolve, France) were added to both eluents. Ionization and data acquisition with data-dependent MS² scans (top 15) was performed as described previously ^37,38^. Lipid species were identified according to exact mass, isotopic pattern, retention time and fragmentation ions using Compound Discoverer 3.3 (Thermo Fisher Scientific) and two in silico databases ^39^. The area under the peak was normalized to the internal standard and the protein content. An extraction blank and quality control samples were used to ensure data quality and linearity.

### Differentially expressed gene (DEG) analysis

Global DEGs analysis was performed independently of sampling time. DESeq2 ^40^ with an adjusted p value of 0.05 (Benjamini-Hochberg Procedure) and a log2 fold-change higher or low than zero was considered as upregulated or downregulated genes, respectively.

### Rhythm analyses

To identify diurnal (24-hour) oscillations in gene expression, we combined four independent algorithms: CircaN ^41^, JTK cycle ^42^, Metacycle ^43^, and DryR ^44^. A single list of rhythmic genes was input in CircaCompare ^16^ to compare rhythmic parameters between groups. For mesor and amplitude comparison, CircaCompare was allowed to fit a sine curve irrespective of meeting rhythmicity thresholds. Phase comparison was only performed in robustly rhythmic genes as described ^11^. CircaN, JTK cycle, and Metacycle algorithms were run in CircaN algorithm using the default parameters. An exact period of 24 h was used. The significance of rhythmicity was set as p < 0.01 for each method. For all comparisons in CircaCompare a p value of 0.05 was used. The same pipeline was done for lipidomics samples, except DryR method was not included. For heatmap and rose plots, phase estimation was extracted from CircaCompare. Rhythmic gene visualization was made using CircaCompare algorithm or ggplot (method “lm” and formula = y ∼ 1 + sin(2*pi*x/24) + cos(2*pi*x/24). Methodological constraints led us to perform differential rhythm analysis using CircaCompare pairwise, without adjusting for multiple comparisons, potentially inflating the rate of false-positive results.

### Gene (GSEA) and phase set enrichment analysis (PSEA)

Enrichment analyses were performed using Gene Ontology (GO) annotations for Biological Processes with DAVID 6.8 software ^45^ and cut-offs of 3 genes per process and a p-value < 0.05. REVIGO (reduction of 0.7 and default conditions) was used to reduce redundant calls in GSEA ^46^. The phases of rhythmic genes were estimated using CircaSingle, round up to exact number, and used for PSEA analysis ^15^. Mouse gene names were converted to human with the assistance of geneName (version 0.2.3). A minimum of 10 genes and a p value of < 0.01 were selected. Biological processes IDs were extracted using the biological processes’ names using the version 7.5.1 (http://geneontology.org/docs/download-ontology/). Processes were filtered in REVIGO (0.7 reduction). The phase of biological processes was estimated by calculating the median from all genes belonging to a biological process. To facilitate comprehension, enriched processes from each genotype were merged into a single list, manually curated by assigning each biological process to a higher hierarchical biological process class. The difference was performed by subtracting the phase from the groups.

### *In vitro* experiments

AML-12 cells, sourced from the ATCC Biobank (catalog no. CRL-2254), were cultured in Gibco Dulbecco’s Modified Eagle Medium supplemented with Nutrient Mixture F12 (DMEM/F12, ThermoFisher). The growth medium was enriched with 1% penicillin-streptomycin, 1% insulin-transferrin-selenium (ITS, ThermoFisher), and 10% non-heat-inactivated fetal bovine serum (FBS, ThermoFisher). Additionally, 10 nM dexamethasone (Sigma-Aldrich) was included to maintain the cells, which were incubated at 37 °C in a humidified atmosphere with 5% CO2. For specific experiments, dexamethasone was excluded, and regular FBS was substituted with charcoal/dextrane-treated FBS (Hyclone) to reduce T_4_ and T_3_ concentrations by 40% and 90%, respectively, in addition to reducing endogenous hormones.

Initially, 10^5^ AML-12 cells were seeded per well in 12-well plates. After 24 hours, the cells underwent synchronization with 200 nM dexamethasone for a duration of 2 hours. Post-synchronization, cells were washed with PBS, and drug pre-treatments were administered for 48h. Subsequent to the drug pre-treatments, a 0.25 mM concentration of BSA-palmitate saturated fatty acid complex (5 mM, 7:1, Cayman Chemical) was introduced. At 48 hours post-palmitate addition, both cells and supernatant were collected. The collection protocol involved centrifugation and washing steps, followed by incubation with Zombie NIR dye (1:500 dilution, Biolegend), Bodipy-Cholesterol (1 µM, MedChem Express), and adipored (0.3 µL per mL, Lonza) at room temperature. Each incubation step was combined with centrifugation and washing. Cells were then fixed in 4% paraformaldehyde (PFA) for 30 minutes at room temperature, subsequently stored for fluorescence-activated cell sorting (FACS) analysis. All drugs were dissolved in dimethyl sulfoxide (DMSO), ensuring the final DMSO concentration remained below 1% in all treatment conditions. The resmetiron (MedchemExpress) concentration was determined based on known EC50 (0.2 µM). T_3_ concentration was 100 nM.

Samples were assessed in Cytek Aurora at the Cell Analysis Core Facility (CAnaCore facility at the University of Lübeck). Data were acquired using predefined templates with adjustments for optimal event rates and detector gains. Spectral unmixing was applied using the manufacturer’s software with single-stained controls to resolve fluorescent populations. At least 10,000 events were captured for each sample. Gating strategies were implemented using FlowJO V10 software, with each gate clearly defined in terms of fluorescence. Data was analyzed using Two-Way ANOVA followed by false discovery rate (Two-stage linear step-up procedure of Benjamini, Krieger, and Yekutieli) correction for multiple comparisons in Prisma (Version 10).

## DATA AVAILABILITY

Transcriptome and lipidomics data are being deposited in Gene Expression Omnibus (GEO) and Metabolomics Workbench, respectively, and will be fully available upon publication. Microarrays from low- and high-TH state mice were obtained from GSE199998.

## ACKNOWLEDGEMENTS AND FUNDING

This work was supported by grants of the German Research Foundation (DFG) to HO 353-10/1, GRK-1957, and CRC/TR 296 “LOCOTACT” (ID 424957847, TP13 and TP14). The authors thank Dr. Sven Geisler for support and fruitful advice in the flow cytometry experiments.

---

**Figure S1:**
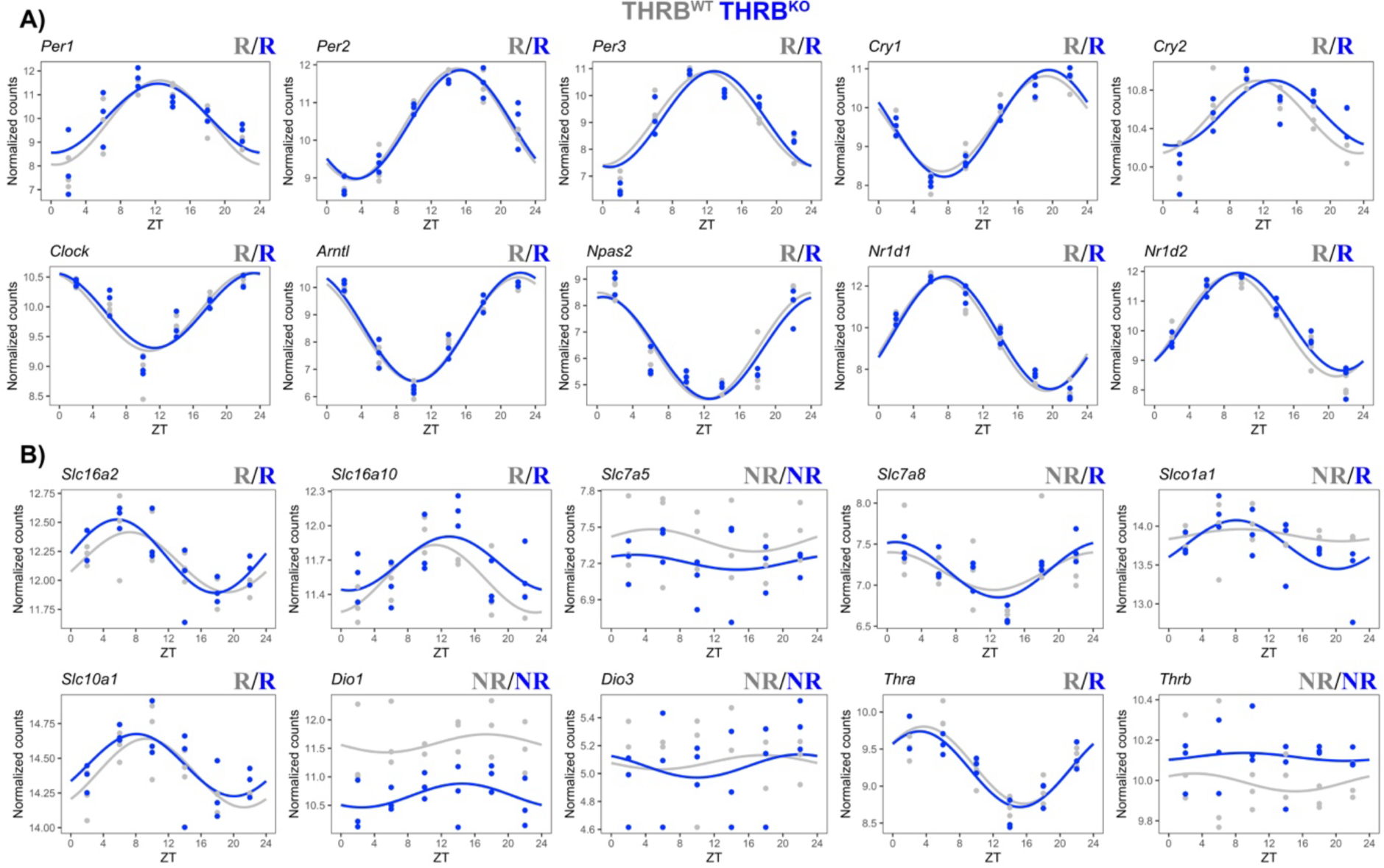
Evaluation of core clock gene and thyroid hormone (TH) modulators in THRB^WT^ compared to THRB^KO^. A) Core clock genes are shown. B) TH modulators, transporters, DIOs, and receptors are shown. Rhythm analysis performed by CircaCompare. N = 3 independent samples per ZT and condition.

**Figure S2:**
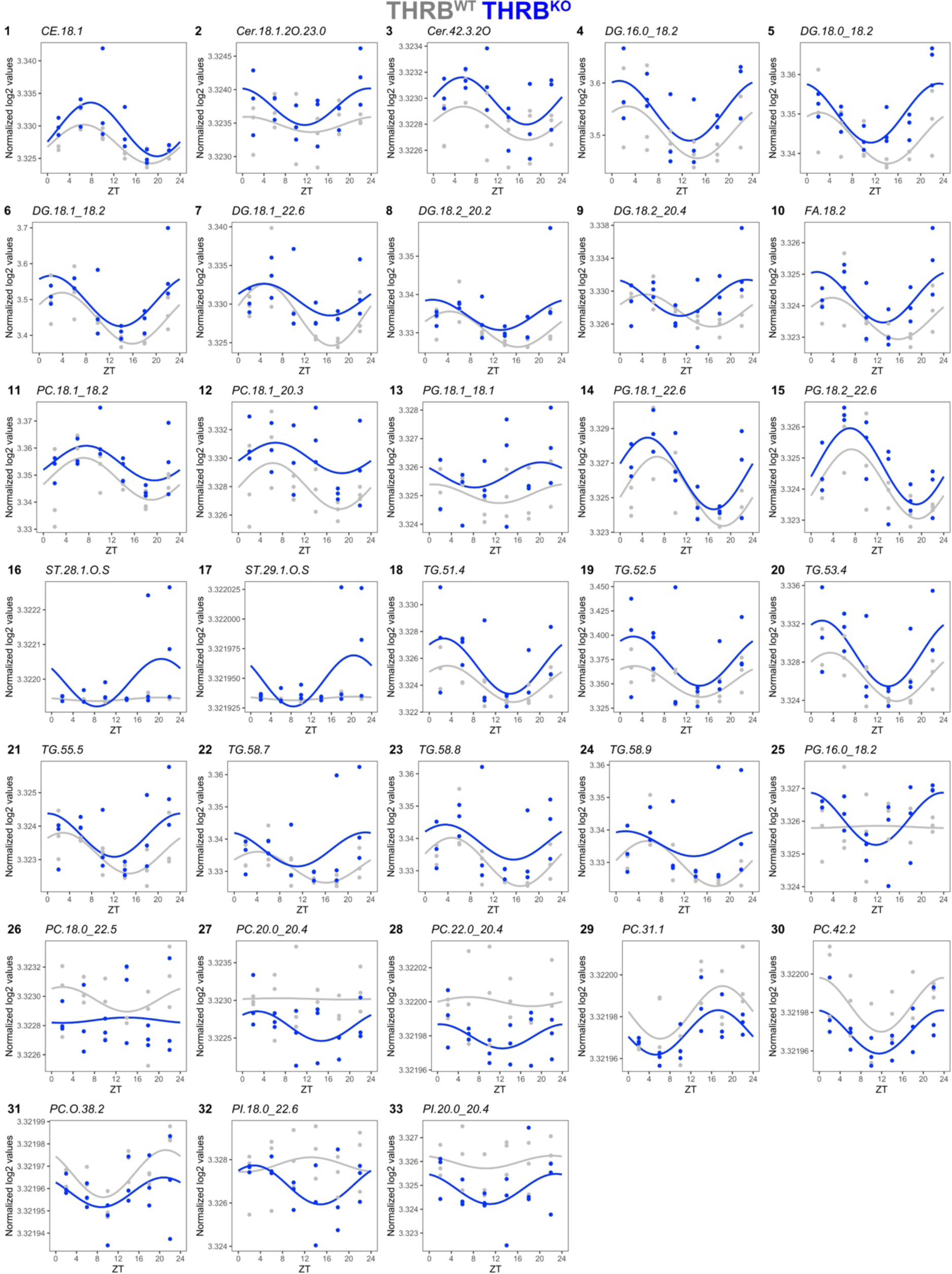
Rewiring of liver lipidome in the absence of THRB. All lipids that show rhythm parameter alteration are depicted. Rhythm analysis performed by CircaCompare. N = 3 independent samples per ZT and condition.

**Figure S3:**
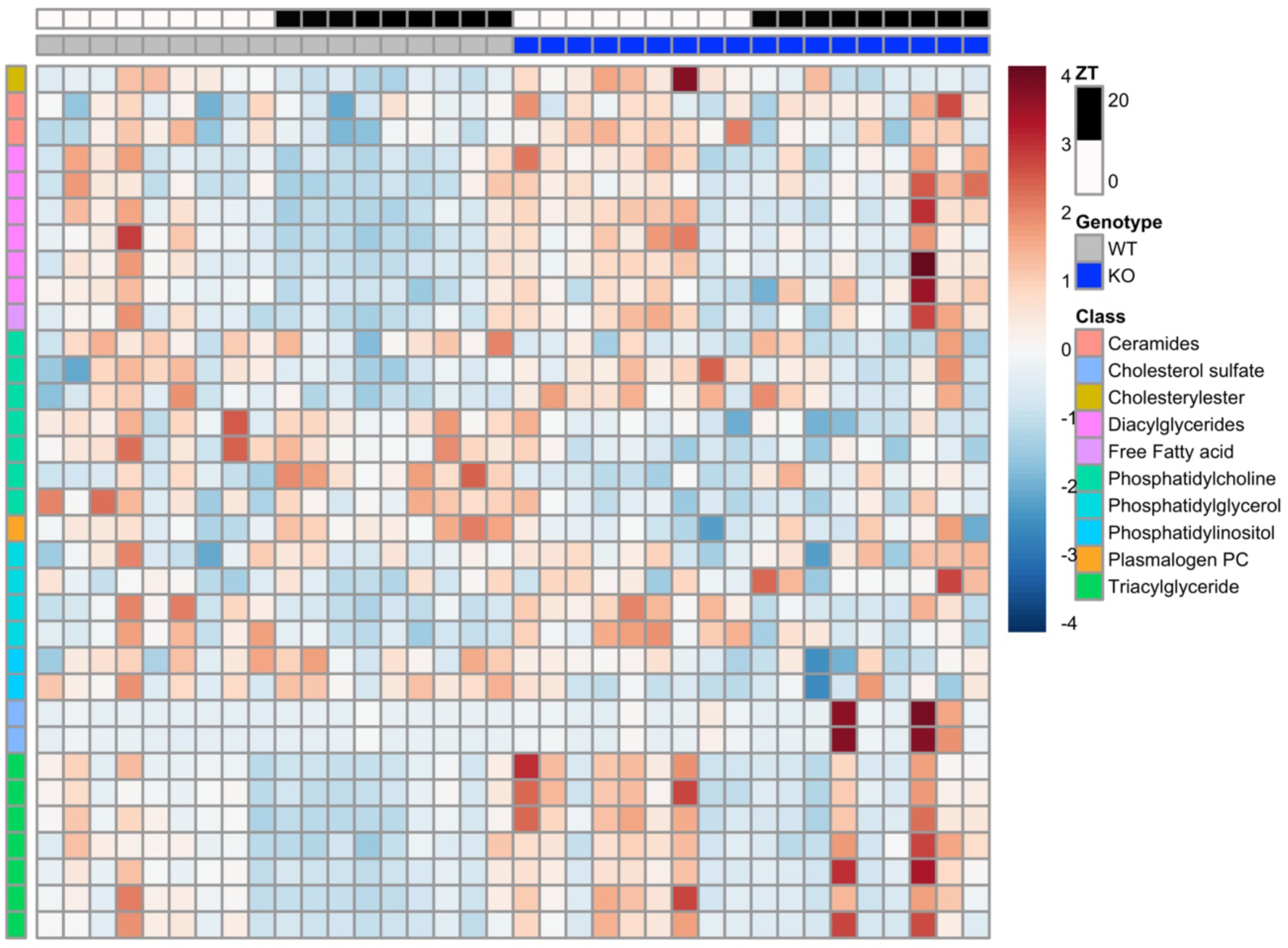
Visualization of the rewiring of liver lipidome in the absence of THRB. Heatmaps show all lipids that show rhythm parameter alteration. Rhythm analysis performed by CircaCompare. N = 3 independent samples per ZT and condition.

**Figure S4:**
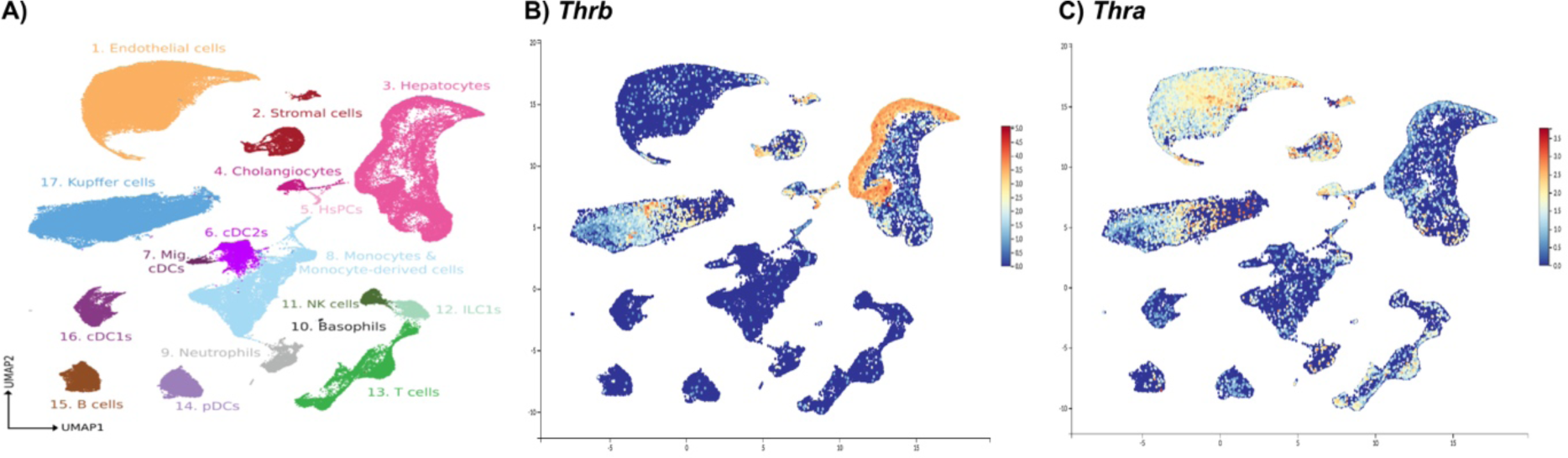
Exploration of THRA and THRB signal in liver cells using public single-cell RNAseq database. A) Liver Cell Atlas (https://www.livercellatlas.org/index.php) was accessed and natural log normalized gene expression was interrogated. Parameters were the following: no cut-off and order dots were used.

## REFERENCES

1. Wirth, E. K., Puengel, T., Spranger, J. & Tacke, F. Thyroid hormones as a disease modifier and therapeutic target in nonalcoholic steatohepatitis. Expert Review of Endocrinology & Metabolism 17, 425–434 (2022).

2. Groeneweg, S., van Geest, F. S., Peeters, R. P., Heuer, H. & Visser, W. E. Thyroid Hormone Transporters. Endocrine Reviews 41, 146–201 (2020).

3. Luongo, C., Dentice, M. & Salvatore, D. Deiodinases and their intricate role in thyroid hormone homeostasis. Nat Rev Endocrinol 15, 479–488 (2019).

4. Mullur, R., Liu, Y.-Y. & Brent, G. A. Thyroid hormone regulation of metabolism. Physiological reviews 94, 355–382 (2014).

5. Sinha, R. A., Singh, B. K. & Yen, P. M. Direct effects of thyroid hormones on hepatic lipid metabolism. Nature Reviews Endocrinology 14, 259–269 (2018).

6. Sinha, R. A., Singh, B. K. & Yen, P. M. Thyroid hormone regulation of hepatic lipid and carbohydrate metabolism. Trends Endocrinol Metab 25, 538–45 (2014).

7. de Assis, L. V. M. & Oster, H. The circadian clock and metabolic homeostasis: entangled networks. Cellular and molecular life sciences : CMLS (2021) doi:10.1007/s00018-021-03800-2.

8. Takahashi, J. S. Transcriptional architecture of the mammalian circadian clock. Nature reviews. Genetics 18, 164–179 (2017).

9. Oster, H. et al. The Functional and Clinical Significance of the 24-Hour Rhythm of Circulating Glucocorticoids. Endocrine Reviews 38, 3–45 (2017).

10. Gamble, K. L., Berry, R., Frank, S. J. & Young, M. E. Circadian clock control of endocrine factors. Nat Rev Endocrinol 10, 466–75 (2014).

11. de Assis, L. V. M. et al. Rewiring of liver diurnal transcriptome rhythms by triiodothyronine (T3) supplementation. eLife 11, (2022).

12. Forrest, D., Erway, L. C., Ng, L., Altschuler, R. & Curran, T. Thyroid hormone receptor β is essential for development of auditory function. Nat Genet 13, 354–357 (1996).

13. de Assis, L. V. M. et al. Tuning of liver circadian transcriptome rhythms by thyroid hormone state in male mice. Sci Rep 14, 640 (2024).

14. de Assis, L. V. M., Demir, M. & Oster, H. Nonalcoholic Steatohepatitis Disrupts Diurnal Liver Transcriptome Rhythms in Mice. Cellular and Molecular Gastroenterology and Hepatology 16, 341–354 (2023).

15. Zhang, R., Podtelezhnikov, A. A., Hogenesch, J. B. & Anafi, R. C. Discovering Biology in Periodic Data through Phase Set Enrichment Analysis (PSEA). J Biol Rhythms 31, 244–257 (2016).

16. Parsons, R., Parsons, R., Garner, N., Oster, H. & Rawashdeh, O. CircaCompare: a method to estimate and statistically support differences in mesor, amplitude and phase, between circadian rhythms. Bioinformatics 36, 1208–1212 (2020).

17. Kannt, A. et al. Activation of thyroid hormone receptor-β improved disease activity and metabolism independent of body weight in a mouse model of non-alcoholic steatohepatitis and fibrosis. British Journal of Pharmacology 178, 2412–2423 (2021).

18. Wenzek, C. et al. The interplay of thyroid hormones and the immune system – where we stand and why we need to know about it. Eur J Endocrinol 186, R65–R77 (2022).

19. Guilliams, M. et al. Spatial proteogenomics reveals distinct and evolutionarily conserved hepatic macrophage niches. Cell 185, 379–396.e38 (2022).

20. de Assis, L. V. M. de et al. Tuning of liver circadian transcriptome rhythms by thyroid hormone state in male mice. Scientific Reports (2024).

21. Adamovich, Y. et al. Circadian clocks and feeding time regulate the oscillations and levels of hepatic triglycerides. Cell metabolism 19, 319–30 (2014).

22. Sprenger, R. R. et al. Lipid molecular timeline profiling reveals diurnal crosstalk between the liver and circulation. Cell Reports 34, 108710 (2021).

23. Männistö, V. et al. Total liver phosphatidylcholine content associates with non-alcoholic steatohepatitis and glycine N-methyltransferase expression. Liver Int 39, 1895–1905 (2019).

24. van der Veen, J. N. et al. The critical role of phosphatidylcholine and phosphatidylethanolamine metabolism in health and disease. Biochimica et Biophysica Acta (BBA) - Biomembranes 1859, 1558–1572 (2017).

25. Arendt, B. M. et al. Nonalcoholic fatty liver disease is associated with lower hepatic and erythrocyte ratios of phosphatidylcholine to phosphatidylethanolamine. Appl. Physiol. Nutr. Metab. 38, 334–340 (2013).

26. Li, Z. et al. The ratio of phosphatidylcholine to phosphatidylethanolamine influences membrane integrity and steatohepatitis. Cell Metabolism 3, 321–331 (2006).

27. de Assis, L. V. M., Demir, M. & Oster, H. The role of the circadian clock in the development, progression, and treatment of non-alcoholic fatty liver disease. Acta Physiologica 237, e13915 (2023).

28. Petersen, M. C. & Shulman, G. I. Roles of Diacylglycerols and Ceramides in Hepatic Insulin Resistance. Trends in Pharmacological Sciences 38, 649–665 (2017).

29. Chaurasia, B. et al. Targeting a ceramide double bond improves insulin resistance and hepatic steatosis. Science 365, 386–392 (2019).

30. Lyu, K. et al. A Membrane-Bound Diacylglycerol Species Induces PKCɛ-Mediated Hepatic Insulin Resistance. Cell Metabolism 32, 654–664.e5 (2020).

31. Araki, O., Ying, H., Zhu, X. G., Willingham, M. C. & Cheng, S. Y. Distinct dysregulation of lipid metabolism by unliganded thyroid hormone receptor isoforms. Molecular endocrinology 23, 308–15 (2009).

32. Hu, L. et al. Discovery of Highly Potent and Selective Thyroid Hormone Receptor β Agonists for the Treatment of Nonalcoholic Steatohepatitis. J. Med. Chem. 66, 3284–3300 (2023).

33. Vatner, D. F. et al. Thyroid hormone receptor-β agonists prevent hepatic steatosis in fat-fed rats but impair insulin sensitivity via discrete pathways. Am J Physiol Endocrinol Metab 305, E89–100 (2013).

34. Erion, M. D. et al. Targeting thyroid hormone receptor-β agonists to the liver reduces cholesterol and triglycerides and improves the therapeutic index. Proceedings of the National Academy of Sciences 104, 15490–15495 (2007).

35. Lopez-Alcantara, N., Oelkrug, R., Sentis, S. C., Kirchner, H. & Mittag, J. Lack of thyroid hormone receptor beta is not detrimental for non-alcoholic steatohepatitis progression. iScience 26, 108064 (2023).

36. Bruinstroop, E. et al. Early induction of hepatic deiodinase type 1 inhibits hepatosteatosis during NAFLD progression. Mol Metab 53, 101266 (2021).

37. Aherrahrou, R. et al. CYP17A1 deficient XY mice display susceptibility to atherosclerosis, altered lipidomic profile and atypical sex development. Sci Rep 10, 8792 (2020).

38. Karsai, G. et al. DEGS1-associated aberrant sphingolipid metabolism impairs nervous system function in humans. J Clin Invest 129, 1229–1239 (2019).

39. Kind, T. et al. LipidBlast in silico tandem mass spectrometry database for lipid identification. Nat Methods 10, 755–758 (2013).

40. Love, M. I., Huber, W. & Anders, S. Moderated estimation of fold change and dispersion for RNA-seq data with DESeq2. Genome Biol 15, 550 (2014).

41. Rubio-Ponce, A., et al. Combined statistical modeling enables accurate mining of circadian transcription. NAR Genomics and Bioinformatics 3, (2021).

42. Hughes, M. E., Hogenesch, J. B. & Kornacker, K. JTK_CYCLE: an efficient nonparametric algorithm for detecting rhythmic components in genome-scale data sets. Journal of biological rhythms 25, 372–380 (2010).

43. Wu, G., Anafi, R. C., Hughes, M. E., Kornacker, K. & Hogenesch, J. B. MetaCycle: an integrated R package to evaluate periodicity in large scale data. Bioinformatics 32, 3351–3353 (2016).

44. Weger, B. D. et al. Systematic analysis of differential rhythmic liver gene expression mediated by the circadian clock and feeding rhythms. Proceedings of the National Academy of Sciences of the United States of America 118, e2015803118 (2021).

45. Huang, D. W., Sherman, B. T. & Lempicki, R. A. Systematic and integrative analysis of large gene lists using DAVID bioinformatics resources. Nature protocols 4, 44–57 (2009).

46. Supek, F., Bošnjak, M., Škunca, N. & Šmuc, T. REVIGO summarizes and visualizes long lists of gene ontology terms. PloS one 6, e21800 (2011).

